# Evoking the N400 Event-Related Potential (ERP) Component Using a Publicly Available Novel Set of Sentences with Semantically Incongruent or Congruent Eggplants (Endings)

**DOI:** 10.1101/2022.03.04.483039

**Authors:** Kathryn K. Toffolo, Edward G. Freedman, John J. Foxe

## Abstract

During speech comprehension, the ongoing context of a sentence is used to predict sentence outcome by limiting subsequent word likelihood. Neurophysiologically, violations of context-dependent predictions result in amplitude modulations of the N400 event-related potential (ERP) component. While N400 is widely used to measure semantic processing and integration, no publicly-available auditory stimulus set is available to standardize approaches across the field. Here, we developed an auditory stimulus set of 404 sentences that utilized the semantic anomaly paradigm, provided cloze probability for all stimuli, and was developed for both children and adults. With 20 neurotypical adults, we validated that this set elicits robust N400’s, as well as two additional semantically-related ERP components: the late positivity component and the recognition potential. This stimulus set and the 20 high-density (128-channel) electrophysiological datasets are made publicly available here to promote data sharing and reuse. Future studies that use this stimulus set to investigate sentential semantic comprehension in both control and clinical populations may benefit from the increased comparability and reproducibility within this field of research.

## 1. Introduction

The ability to use prior context to predict a particular sentence ending is a key feature of fluid language comprehension ^1^. In essence, sentence outcome can be predicted because the words already heard limit the words that can sensibly terminate the sentence. The relationship between prior context and the target word is quantified by ‘cloze probability’ (CP) ^2,3^, which is a measure of how well a predicted word matches the context. For example, in the sentence ‘*Finally, the climbers reached the top of the ---*’ ^4^, one might reasonably predict that the final word is ‘m*ountain*’ or ‘*hill*’. In this example, mountain has a high association with and is semantically congruent with the prior sentence context and would have a high CP. ‘Hill’ is also a reasonable ending to the sentence, but perhaps a little less likely, and as such would have a lower CP. Conversely, the word ‘tulip’ would be a very unexpected ending to this sentence, is semantically incongruent and would be associated with an extremely low CP. Prior analysis of electroencephalography (EEG) has identified a broad negative event related potential (ERP) that peaks around 400 ms after the presentation of semantically related or unrelated stimuli ^2,3,5–7^. This N400 ERP component is evoked by both congruent and incongruent endings, but when incongruent stimuli are presented, the amplitude of the N400 component is substantially increased ^2,3,5–8^. Processing of congruent stimuli is postulated to be easier since it follows the contextual narrowing of expectations, whereas incongruent stimuli result in larger N400 amplitudes due to the increased difficulty of integrating unexpected words ^3,5,9–16^. This N400 ERP component is an established index of semantic processing and integration and has proven a very valuable research tool ^2,5,9,11,17,18^.

Two other ERP components reported to be sensitive to semantic congruence are the recognition potential (RP) and the late positivity component (LPC). The RP is an N2-like component ^19–21^ that occurs over parietal-occipital regions ^17,18^ and is thought to be generated in the left fusiform gyrus or visual word form area (VMFA) ^19,22^. The RP shows larger amplitudes in response to congruent stimuli ^20,23^ and as such, has been generally related to expectancy ^19^ and contextual semantic processing ^22^. More specifically, the RP has been suggested to represent the point at which an individual recognizes the target word following congruent context ^21^. The terms LPC and P600 have tended to be used interchangeably in the literature, with LPC being used most often when the work is specifically addressing responses to semantic contexts ^6,18,32,24–31^.The LPC has a more positive amplitude in response to incongruent stimuli ^26^ and has been related to semantic difficulty or modulations ^18,24–26,32^. The LPC is thought to reflect the reanalysis of semantic information because it occurs after the N400 ^7,27,30,33^. Studies that have used the P600 nomenclature have shown that it is also sensitive to syntactic context ^7,18,24,25,30,34–36^, whereas it has been argued that the amplitude of the LPC is related to the integration of both semantic and syntactic information ^10,18,24,26,29,32,34,37,38^. There is no consensus on the interpretation of the P600 ERP as a whole, nor a consensus on its topographic distribution ^18,25,27,31,32,35,39^.

Semantic studies have used various task designs to understand the nuanced linguistic sensitivities of the N400 and LPC. This has resulted in task dependent topographic differences and various interpretations of the relationship that these components have with prediction, semantic processing, and language comprehension ^5,18,42,43,27,29–32,39–41^. It goes without saying that it is important to use diverse methodologies to understand the linguistic sensitivity of these components. However, the lack of a standardized task design can result in conflicting conclusions about the extent of one’s semantic ability, especially when investigating atypical populations such as Autism Spectrum Disorder (ASD) ^4,7,46–50,18,24–27,33,44,45^. Although a CP validated, lexical N400 stimulus set for adults is publicly available ^8^, to our knowledge, a CP validated, auditory and lexical N400 stimulus set meant for children and adults does not exist. The purpose of this study was to create a stimulus set that targeted auditory, sentential semantic comprehension using a semantic anomaly N400 task design. This stimulus set measured the predictability or CP of each sentence, was appropriate for both children and adults, and could be used to investigate both the auditory and lexical semantic processing of neurotypical (NT) populations and populations known for disorders that impact language and communication skills, such as ASD. This stimulus set and the current dataset are made publicly available. Future semantic studies that use this stimulus set may benefit from the increased comparability and reproducibility within this field of research.

## 2. Results

### 2.1. Accuracy and midline N400 ERP

All participants were able to correctly identify congruent and incongruent endings with at least 97% accuracy. All responses were included in final analyses. Previous studies of the N400 response focused on midline electrode locations ^1,4,5^. To illustrate the N400 along the midline evoked by the current stimulus set, the grand mean ERPs (+/− SEM) for 20 participants, from electrode locations (Pz, Cz, and Fz), were characterized during sentences with congruent (blue) and incongruent (red) endings (Figure. 1). Data were aligned to the onset of the last word in each sentence (t = 0) and baselined during the 200 ms preceding this last word. As shown in Figure 1, during congruent sentence endings, there was a positive-going potential recorded at Pz (Figure. 1a) that increased near the onset of the final word. A similar, increasing, positive potential was also seen at the central electrode site Cz (Figure. 1b). In contrast, during sentences that had incongruent endings, there was a marked negative potential that persisted for ~400 ms. At frontal electrode site (Fz), the negative deflection of the ERP after incongruent sentence endings began later than at the more posterior sites (Figure. 1c). Here the ERPs in response to congruent and incongruent sentences diverged at approximately 550 ms after final word presentation. This later negativity during incongruent endings persisted for ~250 ms. Individual response variation is illustrated to the right of each ERP panel, where individual mean responses over 395 - 405 ms after final word onset (gray band in each panel) are shown for congruent (Con) and incongruent (Incon) sentences. For both parietal and central electrodes, the majority of participants clearly had more negative N400 responses during incongruent sentences, but there was individual variation. Four of the 20 subjects had little or no increase in the N400 response. Individual responses were even more variable across subjects at the frontal electrode location (Figure. 1c). For each electrode, a repeated measures ANOVA, which assessed for an effect of condition (congruent vs. incongruent endings) at 400 ms, is detailed in Table 2. There was a main effect of condition at 400 ms for midline electrodes (Cz) and (Pz), but not for the frontal electrode site (Fz). However, there was a main effect of condition for (Fz) at 700 ms (Table 2).

**Table 1.** Demographic data of the participants. Includes gender, age, handedness, primary language, the number of languages known, and whether the participant was included in the final analysis.

| *Subject* | *Gender* | *Age* | *Handedness* | **Language** |  |  | *Final Dataset* |
| --- | --- | --- | --- | --- | --- | --- | --- |
|  |  |  |  | *Primary/First* | *Known* |  |  |
| 01 | Female | 24 | Right | English | Mono-lingual | 1 | Included |
| 02 | Female | 26 | Left | English | Bi-lingual | 2 | Included |
| 03 | Male | 26 | Left | English | Bi-lingual | 2 | Included |
| 04 | Male | 19 | Right | English | Multi-lingual | 7 | Included |
| 05 | Male | 20 | Right | English | Bi-lingual | 2 | Excluded (fell asleep) |
| 06 | Female | 24 | Right | English | Mono-lingual | 1 | Included |
| 07 | Male | 23 | Right | English | Mono-lingual | 1 | Included |
| 08 | Female | 24 | Right | English | Mono-lingual | 1 | Included |
| 09 | Male | 28 | Right | English | Mono-lingual | 1 | Included |
| 10 | Female | 23 | Right | English | Mono-lingual | 1 | Excluded (excessive noise) |
| 11 | Male | 28 | Left | English | Mono-lingual | 1 | Included |
| 12 | Female | 29 | Right | English | Bi-lingual | 2 | Included |
| 13 | Male | 32 | Right | English | Mono-lingual | 1 | Included |
| 14 | Male | 27 | Right | English | Mono-lingual | 1 | Included |
| 15 | Female | 21 | Right | English | Bi-lingual | 2 | Excluded (excessive noise) |
| 16 | Female | 22 | Right | English | Mono-lingual | 1 | Included |
| 17 | Male | 26 | Right | English | Multi-lingual | 3 | Included |
| 18 | Male | 40 | Left | English | Mono-lingual | 1 | Excluded (excessive moving) |
| 19 | Female | 22 | Right | English | Mono-lingual | 1 | Included |
| 20 | Female | 21 | Right | English | Multi-lingual | 3 | Included |
| 21 | Male | 32 | Right | English | Mono-lingual | 1 | Included |
| 22 | Male | 24 | Right | English | Bi-lingual | 2 | Included |
| 23 | Male | 35 | Right | English | Mono-lingual | 1 | Included |
| 24 | Female | 18 | Right | English | Bi-lingual | 2 | Included |

**Table 2.**
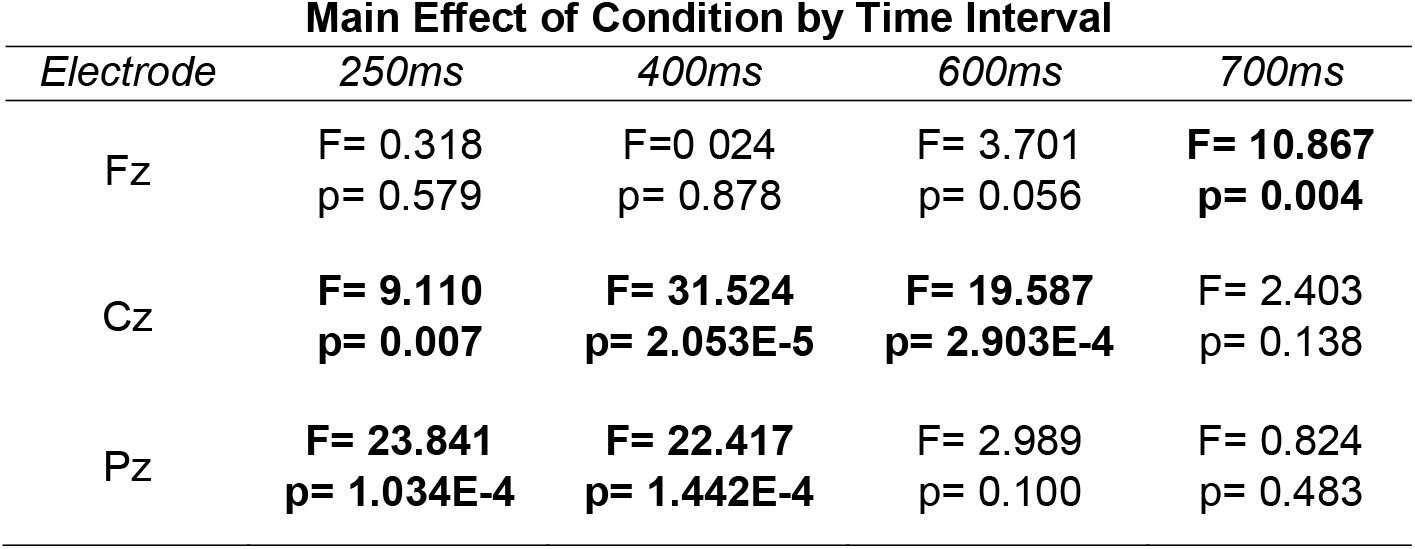
Effect of condition for midline electrodes Fz, Cz, and Pz at four time intervals. Amplitudes for the congruent and incongruent condition were acquired for each electrode, time interval, and participant by averaging the amplitude values from 5 ms before the time point to 5 ms after. Using these values, the effect of condition was assessed using the repeated measure ANOVA. Provided are the F statistic (F) and p-values from this test. Bolded values represent a significant difference between conditions (p < 0.05, two-tailed).

**Figure 1:**
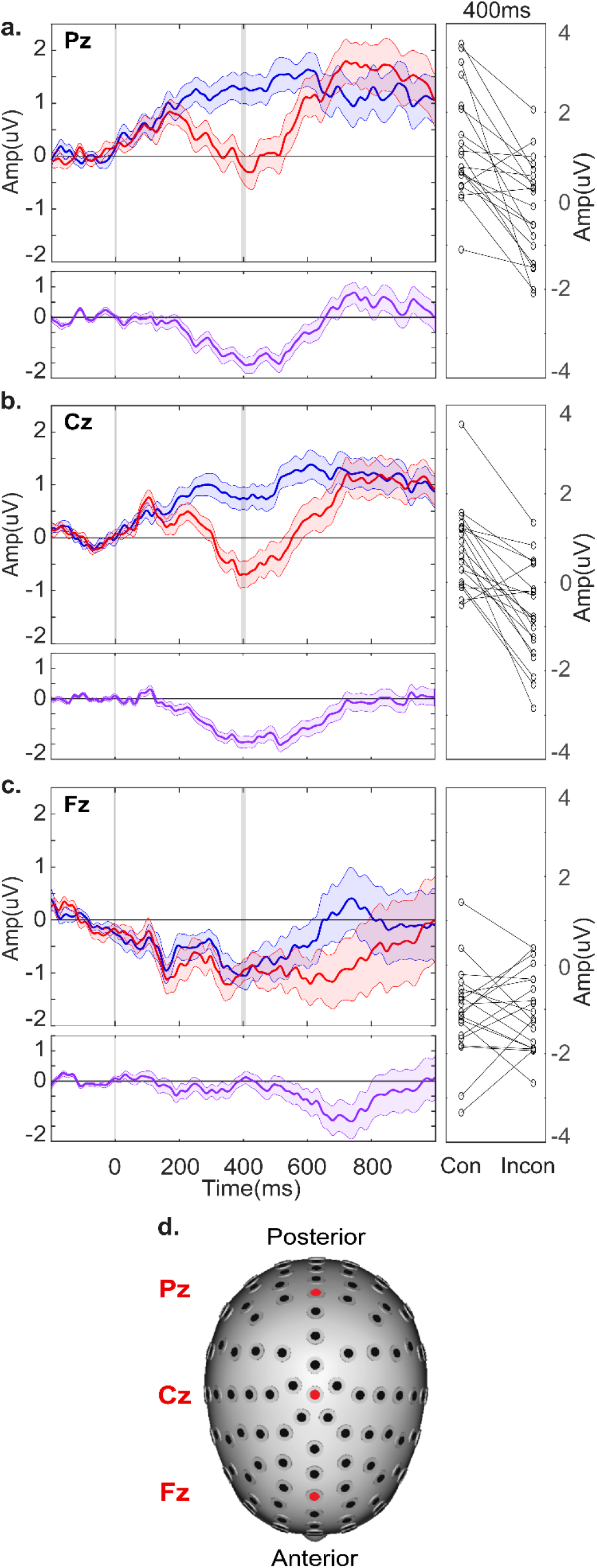
ERP Plots for Midline Electrodes. ERP plots for **a.** Pz, **b.** Cz, and **c.** Fz, show amplitude changes over 1000 ms, per condition (Blue, congruent. Red, Incongruent). Difference plots are in purple (incongruent - congruent amplitudes). The shading around each line represents the s.e.m. To the right of each electrode is the individual participant amplitudes at 400 ms (avg. 395 - 405 ms represented by the gray bar) per condition (congruent vs. incongruent). **d.** Head model showing electrode locations. Statistics for these electrodes are on Table 2.

### 2.2. Topographic representations

In order to identify other regions of interest, the grand mean topography for the 20 included participants was measured over a series of five, 10 ms time windows, aligned to the onset of the last word in each sentence (Figure. 2). The scalp potentials in response to congruent sentence endings are shown in the top row (red = positive; blue = negative). There was clear frontal negativity and central-parietal positivity visible in the 400 ms time window. At later intervals (700 ms and 800 ms), this frontal activity reversed polarity, and the central parietal positivity was more centralized (row ‘a’, columns four and five). Activity across the scalp in response to incongruent sentence endings is shown in row ‘b’. There was a frontal negativity visible in the 400 ms time window that was sustained until 800 ms, where the activity was more positive. The 400 ms time window also displayed bilateral temporal positivities (row ‘b’ of column 2). At later time intervals (600-700 ms), strong positive foci were located over left temporo-parietal and parietal-occipital regions (row ‘b’ of columns three and four). The difference between congruent and incongruent responses is illustrated in row ‘c’. Over the same time windows, row ‘d’ depicts areas of significance (red; p < 0.05, two-tailed). The 400 ms time window of row ‘c’ and ‘d’ illustrates a significantly negative, central-parietal difference between conditions (row ‘c’ and ‘d’ of column 2). Here, there was also significant positivity in fronto-temporal regions of the scalp. Subsequently, positivity was distributed more temporo-parietally and occipitally at later time intervals (600-700 ms) (row ‘c’ and ‘d’ of columns three and four).

**Figure 2:**
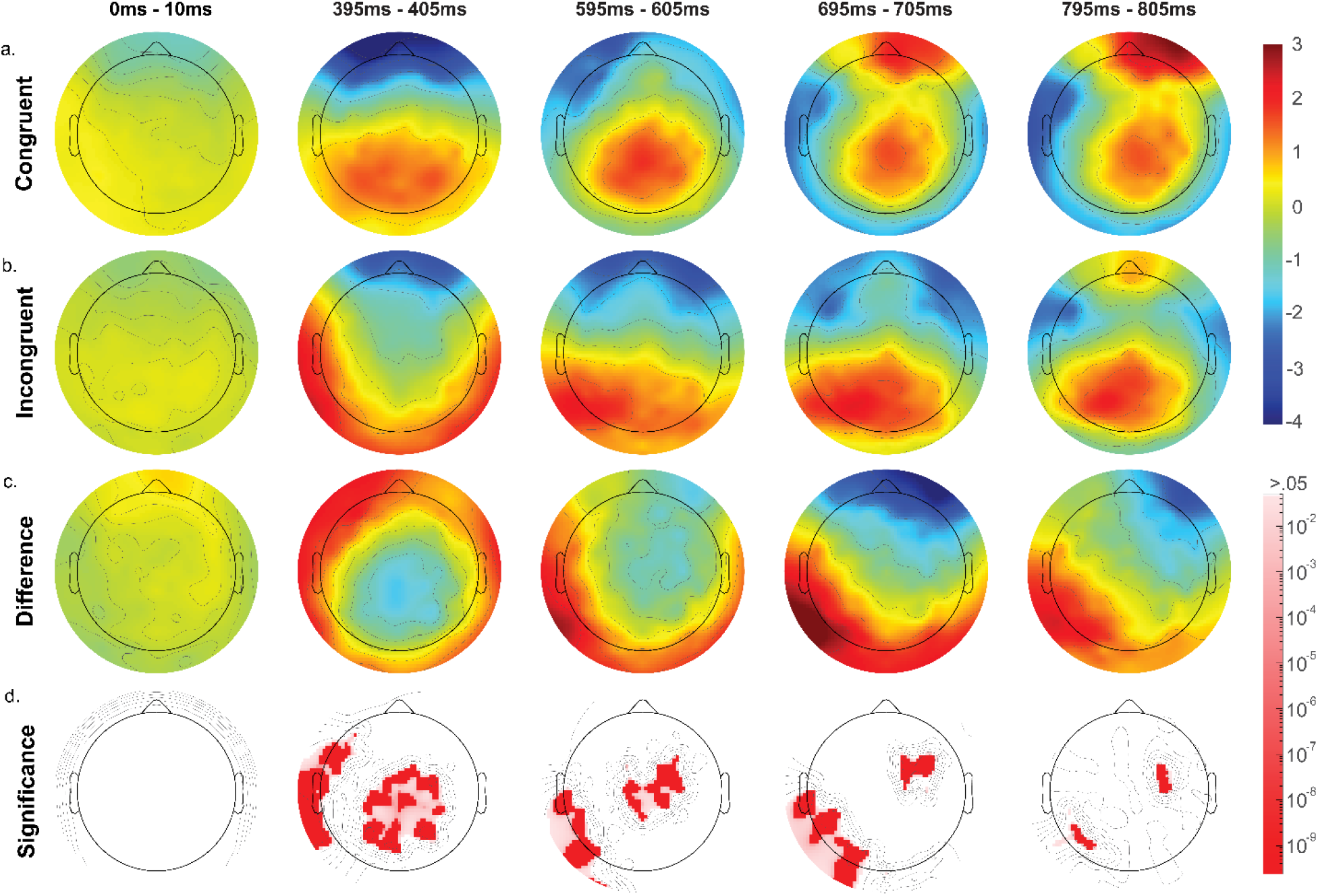
Scalp topography over 5 time points. Distribution of EEG amplitudes based on an average reference. Focused on 5 major time points (0 ms, 400 ms, 600 ms, 700 ms, and 800 ms) averaged over 10 ms. Topographic amplitudes in response to **a.** Congruent stimuli, **b.** Incongruent stimuli. **c.** The difference between conditions (Incongruent - Congruent amplitudes). **d.** T-test cluster plots. White is .05 and greater.

### 2.3. Semantic Sensitivity over left temporal, parietal, and fronto-temporal regions

The grand average ERPs for the left temporal, parietal, and fronto-temporal regions of interest (identified for post hoc analysis via the grand mean topography (Figure. 2)), are characterized in Figures. 3 – 5 respectively. Like midline electrodes (Cz) and (Pz) (Figure. 1a and b), the conditional response at left temporal (T7) and temporo-parietal (TP7) regions peaked 400 ms after final word onset, although with reversed polarity (Figure. 3c and 4a). In response to congruent endings, these regions had a small early negative deflection beginning around 200 ms which persisted for ~200 ms. At the end of this early deflection, ~400 ms after final word onset, there was a substantial negative potential that spanned the remainder of the epoch. This negative potential was mimicked in the incongruent condition following a positive-going potential that began around 150 ms and peaked at approximately at 400 ms (Figure. 3c and 4a). Conditional responses in left parietal (P7) and parietal-occipital (PO7) regions were similar to those from temporo-parietal regions (Figure. 4b and c). However, the onset of divergence between conditional responses began later for more occipital sites (Figure. 4c). An rmANOVA conducted at electrodes (T7) and (P7) showed an effect of condition at 250 ms, 400 ms, 600 ms, and 700 ms (Table 4).

**Table 3.** Alternative main effects for midline electrodes Fz, Cz, and Pz at four time intervals. Amplitudes for the congruent and incongruent condition were acquired for each electrode, effect (i.e. cloze probability (CP), order, language division (LD), and time period (TP) of the experiment), time interval, and participant by averaging the amplitude values from 5 ms before the time point to 5 ms after. Using these values, the main effects of condition and alternative affects were assessed using the repeated measure ANOVA. Provided are the F statistic (F) and p-values from this test. Bolded values represent a significant difference between conditions (p < 0.05, two-tailed). Italicized values represent where the assumption of sphericity (p < 0.05, two-tailed) was violated using the Mauchly’s test of sphericity.

| Main Effect | Time Interval |  |  |  |
| --- | --- | --- | --- | --- |
|  | 250ms | 400ms | 600ms | 700ms |
|  | Electrode Fz |  |  |  |
| Close Probability | F= 1.088 p= 0.361 | F= 0.677 p= 0.570 | F= 0.625 p= 0.602 | F= 1.186 p= 0.323 |
| Order | **F= 4.552 p= 0.046** | F= 0.017 p= 0.898 | F= 0.141 p= 0.712 | **F= 4.443 p= 0.049** |
| Language Division | F= 2.854 p= 0.108 | **F= 4.387 p= 0.050** | F= 1.578 p= 0.224 | F= 0.505 p= 0.486 |
| Time Period | F= 0.369 p= 0.776 | F= 0.120 p= 0.948 | F= 0.183 p= 0.907 | F= 0.229 p= 0.876 |
| Condition*CP | *F= 1.873 p= 0.144* | F= 1.544 p= 0.213 | F= 0.823 p= 0.486 | *F= 1.576 p= 0.205* |
| Condition*Order | F= 0.061 p= 0.807 | F= 0.846 p= 0.369 | F= 1.578 p= 0.224 | F= 0.470 p= 0.501 |
| Condition*LD | F= 0.003 p= 0.959 | F= 0.290 p= 0.597 | F= 0.087 p= 0.771 | F= 0.392 p= 0.539 |
| Condition*TP | F= 0.183 p= 0.907 | F= 0.356 p= 0.785 | *F= 0.568 p= 0.639* | *F= 0.401 p= 0.753* |
| Main Effect | Electrode Cz |  |  |  |
|  | 250ms | 400ms | 600ms | 700ms |
|  | Electrode Cz |  |  |  |
| Close Probability | F= 1.403 p= 0.252 | *F= 1.1665 p= 0.331* | F= 2.288 p= 0.088 | F= 0.101 p= 0.959 |
| Order | F= 0.917 p= 0.250 | F= 0.423 p= 0.523 | F= 0.316 p= 0.518 | F= 0.029 p= 0.866 |
| Language Division | F= 1.364 p= 0.247 | F= 1.247 p= 0.278 | F= 0.102 p= 0.752 | F= 1.811 p= 0.194 |
| Time Frame | F= 0.655 p= 0.583 | F= 0.530 p= 0.663 | F= 0.494 p= 0.688 | F= 0.791 p= 0.504 |
| Condition*CP | F= 1.822 p= 0.153 | F= 1.596 p= 0.200 | F= 1.894 p= 0.141 | F= 1.542 p= 0.213 |
| Condition*Order | F= 0.008 p= 0.931 | F= 0.088 p= 0.770 | F= 1.312 p= 0.266 | F= 2.542 p= 0.127 |
| Condition*LD | F= 0.113 p= 0.740 | F= 0.040 p= 0.844 | F= 0.396 p= 0.537 | F= 1.372 p= 0.256 |
| Condition*TP | F=0.433 p=0.730 | F=0.431 p=0.732 | F=0.471 p=0.704 | F=0.568 p=0.638 |
| Main Effect | Electrode Pz |  |  |  |
|  | 250ms | 400ms | 600ms | 700ms |
|  | Electrode Pz |  |  |  |
| Close Probability | *F= 1.165 p= 0.331* | F= 0.762 p= 0.520 | F= 1.886 p= 0.142 | **F= 5.629 p= 0.002** |
| Order | F= 2.628 p= 0.121 | F= 0.548 p= 0.466 | F= 1.041 p= 0.320 | **F= 6.274 p= 0.022** |
| Language Division | F= 1.868 p= 0.188 | F= 2.320 p= 0.144 | F= 0.137 p= 0.715 | F= 0.259 p= 0.616 |
| Time Period | F= 0.578 p= 0.632 | F= 0.777 p= 0.512 | F= 1.250 p= 0.300 | F= 1.512 p= 0.221 |
| Condition*CP | F= 0.797 p= 0.501 | F= 1.049 p= 0.378 | **F= 2.781 p= 0.049** | F= 2.338 p= 0.083 |
| Condition*Order | F= 2.446 p= 0.134 | F= 1.521 p= 0.232 | **F= 9.046 p= 0.007** | **F= 5.019 p= 0.037** |
| Condition*LD | F= 1.350E-5 p= 0.006 | F= 2.589E-4 p= 0.122 | F= 0.077 p=0.784 | F= 1.619 p= 0.219 |
| Condition*TP | F= 0.913 p= 0.441 | *F= 0.700 p= 0.556* | **F= 2.793 p= 0.048** | **F= 4.544 p= 0.006** |

**Table 4.** Main effects for electrodes F7, T7, and P7 at four time intervals. Amplitudes for the congruent and incongruent condition were acquired for each electrode, effect (i.e. cloze probability (CP), order, language division (LD), and time period (TP) of the experiment), time interval, and participant by averaging the amplitude values from 5 ms before the time point to 5 ms after. Using these values, the main effects of condition and alternative affects were assessed using the repeated measure ANOVA. Provided are the F statistic (F) and p-values from this test. Bolded values represent a significant difference between conditions (p < 0.05, two-tailed). Italicized values represent where the assumption of sphericity (p < 0.05, two-tailed) was violated using the Mauchly’s test of sphericity.

| Main Effect | Time Interval |  |  |  |
| --- | --- | --- | --- | --- |
|  | 250ms | 400ms | 600ms | 700ms |
|  | Electrode F7 |  |  |  |
| Condition | **F= 25.089 p= 7.793E-5** | **F= 24.750 p= 8.408E-5** | F= 2.862 p= 0.107 | F= 0.219 p= 0.645 |
| Close Probability | F= 0.213 p= 0.887 | F= 0.282 p= 0.838 | F= 0.662 p= 0.579 | F= 0.401 p= 0.753 |
| Order | F= 0.038 p= 0.847 | F= 0.024 p= 0.878 | F= 0.024 p= 0.878 | F= 0.882 p= 0.359 |
| Language Division | F= 0.352 p= 0.560 | F= 2.897 p= 0.105 | F= 0.158 p= 0.696 | F= 0.013 p= 0.909 |
| Time Period | F= 0.774 p= 0.514 | F= 1.061 p= 0.373 | **F= 3.645 p= 0.018** | F= 1.591 p= 0.201 |
| Condition*CP | F= 0.868 p= 0.463 | F= 0.797 p= 0.501 | F= 0.673 p= 0.572 | F= 0.203 p= 0.894 |
| Condition*Order | F= 0.590 p= 0.452 | F= 0.250 p= 0.623 | F= 3.293 p= 0.085 | F= 0.953 p= 0.341 |
| Condition*LD | F= 2.040 p= 0.169 | F= 0.510 p= 0.484 | F= 2.000 p= 0.173 | F= 0.057 p= 0.814 |
| Condition*TP | F= 0.039 p= 0.990 | F= 0.153 p= 0.927 | F= 0.503 p= 0.682 | F= 0.363 p= 0.780 |
| Electrode T7 |  |  |  |  |
| Condition | **F= 27.709 p= 4.424E-5** | **F= 29.290 p= 3.194E-5** | **F= 25.564 p= 7.016E-5** | **F=21.852 p= 1.651E-4** |
| Close Probability | F= 0.716 p= 0.546 | *F= 1.093 p= 0.360* | F= 1.099 p= 0.357 | **F= 3.003 p= 0.038** |
| Order | F= 1.268 p= 0.274 | F= 1.995 p= 0.174 | F= 0.039 p= 0.845 | F= 0.122 p= 0.731 |
| Language Division | F= 0.027 p= 0.872 | F= 0.612 p= 0.444 | F= 0.393 p= 0.538 | F= 0.365 p= 0.553 |
| Time Period | F= 1.247 p= 0.301 | F= 1.289 p= 0.287 | F= 0.347 p= 0.791 | F= 0.958 p= 0.419 |
| Condition*CP | *F= 0.955 p=0.420* | *F= 1.251 p= 0.300* | *F= 1.461 p= 0.235* | F= 1.919 p= 0.137 |
| Condition*Order | F= 2.049 p= 0.169 | F= 0.347 p= 0.563 | F= 0.028 p= 0.870 | F= 0.025 p= 0.876 |
| Condition*LD | F= 0.301 p= 0.590 | F= 4.757E-9 p= 1.000 | F= 0.091 p= 0.766 | F= 0.904 p= 0.354 |
| Condition*TP | F= 0.168 p= 0.918 | F= 1.096 p= 0.358 | F= 0.602 p= 0.617 | F= 0.193 p= 0.901 |
| Electrode P7 |  |  |  |  |
| Condition | **F= 4.800 p= 0.042** | **F= 8.796 p= 0.008** | **F= 27.060 p= 5.075E-5** | **F= 43.959 p= 2.418E-5** |
| Close Probability | *F= 0.407 p= 0.748* | F= 0.780 p= 0.510 | **F= 10.003 p= 2.150E-5** | F= 1.797 p= 0.158 |
| Order | F= 0.390 p= 0.540 | F= 0.035 p= 0.853 | F= 0.259 p= 0.062 | F= 1.285 p= 0.271 |
| Language Division | F= 1.118 p= 0.304 | F= 1.870 p= 0.187 | F= 0.024 p= 0.879 | F= 0.201 p= 0.659 |
| Time Period | F= 0.353 p= 0.787 | F= 0.874 p= 0.460 | F= 0.882 p= 0.456 | F= 0.992 p= 0.403 |
| Condition*CP | F= 1.502 p= 0.224 | F= 0.037 p= 0.990 | **F= 6.275 p= 9.864E-4** | **F= 2.410 p= 0.076** |
| Condition*Order | F=0.477 p= 0.498 | F= 1.162 p= 0.295 | F= 1.088 p= 0.310 | F= 0.082 p= 0.777 |
| Condition*LD | F= 0.306 p= 0.587 | F= 2.963 p= 0.101 | F= 0.547 p= 0.468 | F= 0.056 p= 0.816 |
| Condition*TP | *F= 0.214 p= 0.886* | F= 0.884 p= 0.455 | F= 0.855 p= 0.470 | *F= 2.227 p= 0.095* |

**Figure 3:**
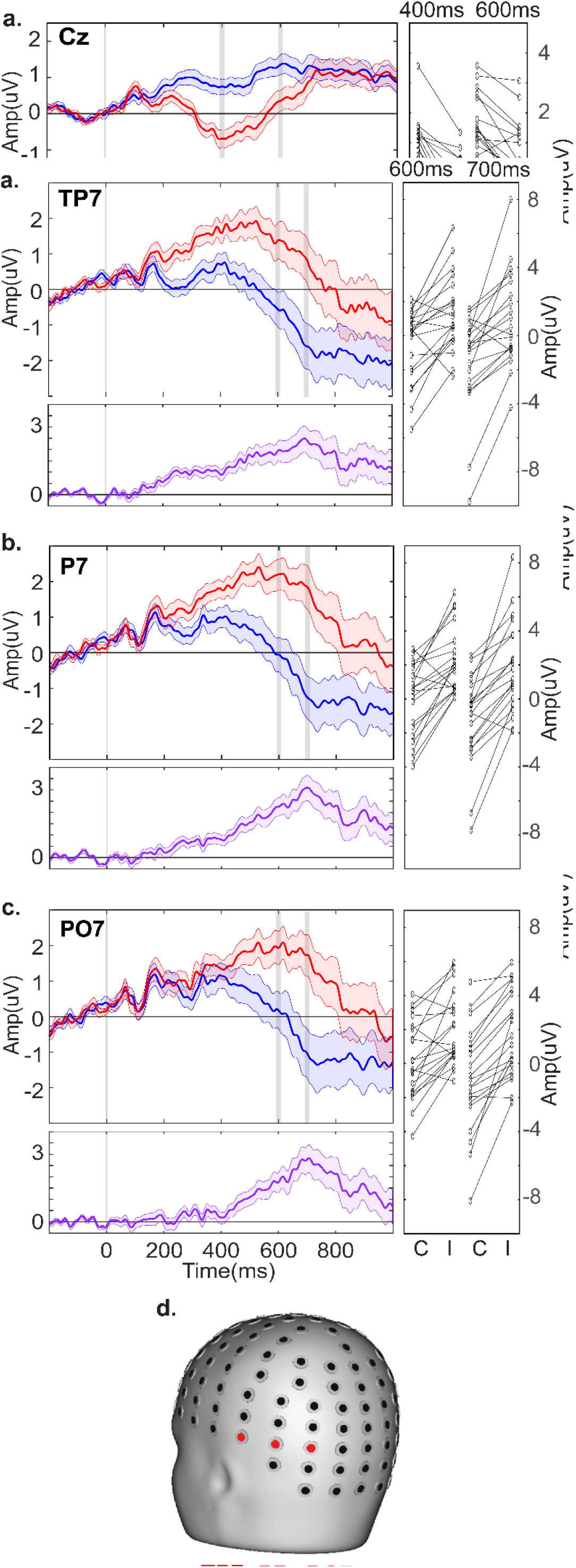
ERP Plots for Temporal Electrodes. ERP plots for **a.** Cz, **b.** C3, and **c.** T7, show amplitude changes over 1000 ms, per condition (Blue, congruent. Red, Incongruent). Difference plots are in purple (incongruent - congruent amplitudes). The shading around each line represents the s.e.m. To the right of each electrode is the individual participant amplitudes at 400 ms and 600 ms (avg. 395 - 405 ms and 595 - 605 ms respectively, represented by the gray bars) per condition (congruent vs. incongruent). **d.** Head model showing electrode locations. Statistics for T7 are on Table 4.

**Figure 4:** ERP Plots for Temporo-Occipital Electrodes. ERP plots for **a.** TP7, **b.** P7, and **c.** PO7, show amplitude changes over 1000 ms, per condition (Blue, congruent. Red, Incongruent). Difference plots are in purple (incongruent - congruent amplitudes). The shading around each line represents the s.e.m. To the right of each electrode is the individual participant amplitudes at 600 ms and 700 ms (avg. 595 - 605 ms and 695 - 705 ms respectively, represented by the gray bars) per condition (congruent vs. incongruent). **d.** Head model showing electrode locations. Statistics for electrode P7 are on Table 4.

**Figure 5:**
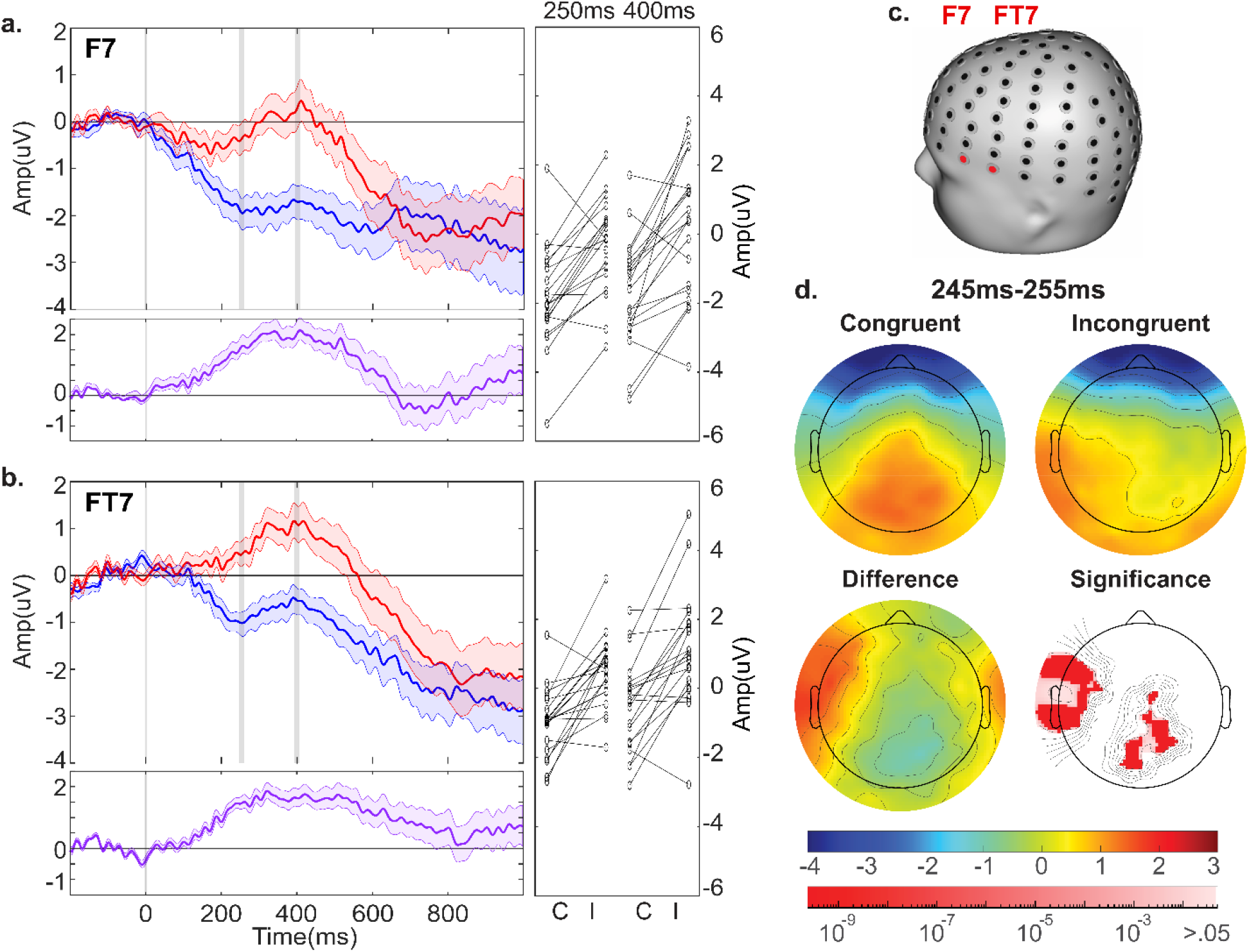
ERP Plots for Electrodes over the Inferior Frontal Gyrus (IFG). ERP plots for **a.** F7 and **b.** FT7, show amplitude changes over 1000 ms, per condition (Blue, congruent. Red, Incongruent). Difference plots are in purple (incongruent - congruent amplitudes). The shading around each line represents the s.e.m. To the right of each electrode is the individual participant amplitudes at 250 ms and 400 ms (avg. 245 - 255 ms and 395 - 405 ms respectively, represented by the gray bars) per condition (congruent vs. incongruent). **d.** Head model showing electrode locations. **e.** Topography averaged between 225 and 250 ms for congruent stimuli, incongruent stimuli, difference between conditions (incongruent - congruent amplitudes), and T-test cluster plots for this time frame (white means the p-value is > .05). Statistics for F7 are on Table 4.

The conditional responses in the left fronto-temporal regions (F7 and FT7) around the inferior frontal gyrus (IFG) (Figure. 5a and b) were similar to those in temporal and temporo-parietal regions (Figure. 3c and 4a). Specifically, in response to incongruent endings, there was a positive going deflection that began ~175 ms after final word onset and peaked at 400 ms, followed by a negative deflection that spanned until 800 ms (Figure. 5a and b). However, in response to congruent sentence endings, fronto-temporal regions had a much more robust early deflection that began ~100 ms and peaked ~250 ms after final word onset. This early peak was then followed by a negative going potential that started at 400 ms and continued for the remainder of the epoch, similar to the congruent response in temporal regions. An rmANOVA conducted at fronto-temporal electrode (F7) showed an effect of condition at both 250 ms and 400 ms (Table 4). In order to more accurately characterize this early peak, the grand mean topography was measured over a 10 ms time window, centered at 250 ms after final word onset (Figure. 5d.). There was substantial frontal negativity in both conditions. In contrast, in response to congruent endings, there was positive activity in central-parietal locations, while in response to incongruent endings, there was a leftward positivity over temporal regions. Both of these central-parietal and left fronto-temporal regions had significant differences between conditions.

### 2.4. Effects of cloze probability

To investigate for an effect of CP on conditional responses, the grand mean ERPs for midline electrodes (Pz, Cz, Fz) were characterized after separating sentences by their cloze probability (CP) score: sentences with 96 - 100% CP (54 sentence pairs; Figure. 6a.); sentences with 90 - <96% CP (56 sentence pairs; Figure. 6b.); sentences with 80 - <90% CP (46 sentence pairs; Figure. 6c.); and sentences with <80% CP (43 sentence pairs; Figure. 6d.). The distribution of CP in this stimulus set is shown in Supplementary Figure 3. Each CP division (Figure. 6a – d) for these electrodes was similar to their respective grand average ERPs depicted in Figure. 1. Furthermore, the response to each condition was similar between all CP divisions. For midline electrodes (Pz) and (Cz), in response to incongruent sentence endings, all divisions had a negative-going potential that began ~200 ms after final word onset, persisted for ~400 ms, and peaked approximately at 400 ms. In response to congruent endings, although the amplitudes varied, all divisions had a positive potential that began at the onset of the final word to ~600 ms. An rmANOVA conducted at four time points (250 ms, 400 ms, 600 ms, and 700 ms) for electrode Cz showed no effect of CP or Condition*CP. However, electrode Pz showed a significant effect of CP at 700 ms, and Condition*CP at 600 ms (Table 3). At the frontal electrode (Fz), regardless of CP division, there was a negative going deflection in response to both congruent and incongruent sentence endings. However, there was no effect of CP or Condition*CP for electrode (Fz) at any time point (Table 3). The effect of CP at frontal, temporal and parietal electrode sites (F7, T7, P7) is characterized in Supplementary Figure 4. There were no effects of CP or CP*Condition at 400 ms for any electrode (Table 4). For parietal electrode (P7), there was an effect of CP at 600 ms and an effect of CP*Condition at both 600 and 700 ms.

**Figure 6:**
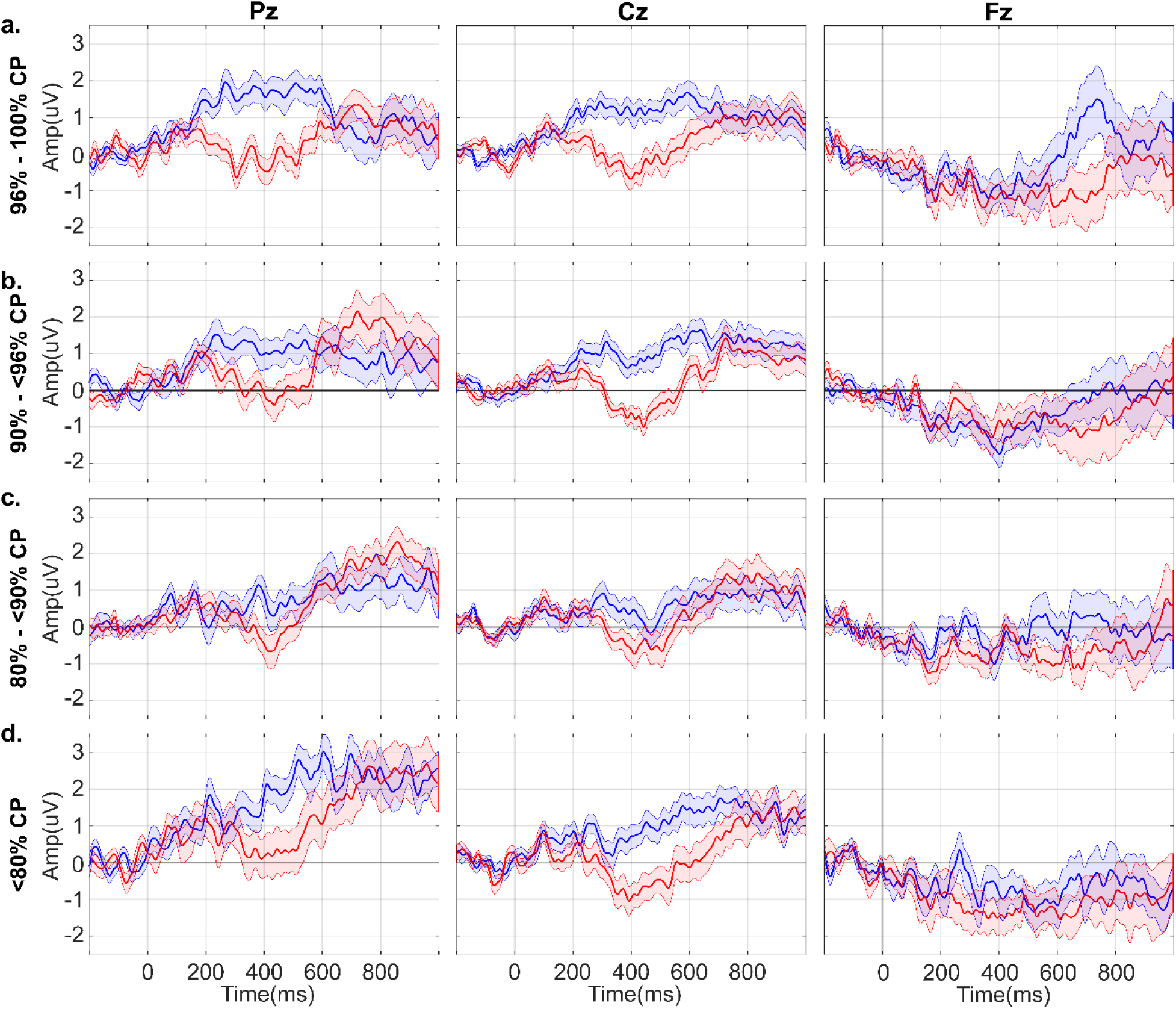
ERP Plots of Midline Electrodes Pz, Cz, and Fz separated by differing levels of Cloze Probability. The ERP plots show amplitude changes over 1000 ms, per condition (Blue, congruent. Red, Incongruent). The shading around each line represents the s.e.m. **a.** Sentences with 96 - 100% cloze probability (54 congruent and 55 incongruent stimuli). **b.** Sentences with 90-<96% cloze probability (57 sentences pairs). **c.** Sentences with 80 - <90% cloze probability (47 sentence pairs). **d.** Sentences with <80% cloze probability (43 congruent and 42 incongruent stimuli). Statistics for these electrodes are on Table 3.

## 3. Discussion

The purpose of this study was to develop and make publicly available, a standardized and validated auditory semantic anomaly stimulus set that evoked the classic N400 ERP component, and to provide a high-density EEG dataset from neurotypical adults (N=20) for use by researchers interested in the neurophysiology of semantic and syntactic processing. This stimulus set did indeed evoke a reliable N400 response across the 20 NT adults in both congruent and incongruent sentences. Consistent with prior literature ^2,3,5–8^, relative to congruent sentence endings, incongruent sentence endings produced a larger amplitude deflection 400 ms after the onset of the final word over central-parietal regions. These data reinforce the interpretation that the N400 represents an index of semantic processing and integration ^2,5,9,11,17,18^. Left-lateralized, semantically-driven responses 400 ms after final word onset were also observed over temporal scalp sites. Significant semantically-related activity was also observed over left anterior regions such as the IFG. There was an early negative deflection peaking 250 - 250 ms post-stimulus onset in response to only congruent sentence endings, consistent with the RP literature ^18–24^. However, the topography of the RP in the prior literature was not over the IFG, but rather over the VMFA ^19,22^. This difference is almost certainly due to the sensory modality in which the stimuli were presented. Unlike some prior studies that used visual words ^19,23^ or pictures ^20^, stimuli here were presented auditorily in order to evoke responses in the auditory cortex and areas commonly associated with language comprehension. These topographic differences suggest that the RP topography is dependent on the sensory modality of the stimuli. There was also significant semantically-related positive activity over temporal and parietal-occipital regions, peaking 600 – 1000 ms post-stimulus onset, that was consistent with LPC (P600) literature ^6,26,33^. We would like to remind the reader that the terms LPC and P600 have been used interchangeably, with LPC being used most often when the work is specifically addressing responses to semantic contexts ^6,18,32,24–31^. For our current purposes, we will refer to this component as “LPC (P600)” in an effort to encompass the terminology of prior studies and avoid confusion.

Because the peak amplitude of the N400 component has been shown to be modulated by expectation ^3,5,9–16^, we investigated the effect of CP on the amplitude of condition specific modulations of the N400. It was hypothesized that as word expectation became less dependent on context (i.e. lower CP), the integration of subsequent words would be more difficult. Data from the current study did not support this hypothesis. For midline, frontal, temporal, and parietal electrode sites, there were no main effects of CP, and there was no CP by Condition interaction at 400 ms. There were however, effects of CP and CP*Condition at later time points over parietal electrode sites, which suggested that the LPC (P600) was more sensitive to CP differences in this stimulus set. This lack of an effect at 400 ms may also be explained by the ‘high’ CP of the majority of our stimuli ^8,51,52^.

An effect of presentation order (i.e. hearing the congruent sentence before its incongruent counterpart, and vice versa) was investigated because the task design included two iterations of each sentence, where the ending was either congruent or incongruent. It was expected that semantic violations of word expectation in newer contexts would be harder to integrate than violations in repeated contexts, but the data did not support this hypothesis (Supplementary Figure 5 and 6; Table 3 and 4). However, there was an effect of Condition*Order at electrode (Pz) that corresponded with more positive amplitudes in response to incongruent sentence endings over the 600 - 900 ms timeframe, when the incongruent pair was presented first (Supplementary Figure 5). This suggests that the LPC (P600) was more sensitive to the presentation order of these stimuli.

Because prior literature has suggested that the LPC (P600) component was sensitive to both semantic and syntactic errors ^10,18,24,26,29,32,34,37,38^, effects of syntactic errors in this stimulus set were also investigated. For this analysis, responses to sentences were separated into 2 linguistic divisions (LD): 1. Sentence pairs with only semantic errors; and 2. Sentence pairs with both semantic and syntactic errors. This analysis found an effect of LD at 400 ms at frontal electrode site Fz. There were no effects of Condition*LD for any electrode at any time point (Supplementary Figure 7 and 8; Table 3 and 4). This suggests that semantic errors were more salient than syntactic errors in this stimulus set, which reinforces that this task was primarily a semantic task for neurotypical adults.

The analyses for an effect of CP, order, and linguistic error had relatively minor effects for this NT sample, which may not be the case for other populations. Future studies that utilize these analyses could provide insight into expectation, memory, or linguistic comprehension differences between NT populations and other populations of interest. Furthermore, semantic studies that make use of this stimulus set may increase the comparability and reproducibility within this field of research.

A number of limitations of this study should be mentioned. First, although inconsistent, there was an effect of Time and Condition*Time over the LPC (P600) (Supplementary Figure 9 and 10; Table 3 and 4). To reduce any time-on-task effects, future studies should follow our break schedule, and consider either enforcing an extended halfway break or reducing the number of stimuli employed. Second, stimuli were presented in a pseudorandomized order. Although the grand average mean for each channel would not be affected by this, pseudorandomization could present bias in other analyses such as an effect of CP, stimulus order, linguistic error, and time. Therefore, differences uncovered here could have been influenced by when in the task the stimuli were presented. To have a more complete analysis of the data, future studies that use this stimulus set should properly randomize the stimuli.

Future research may find that this N400 stimulus set is especially useful for studying the auditory/lexical semantic abilities of atypical populations, such as individuals with ASD, primarily because this stimulus set used a linguistic, semantic anomaly paradigm, which targeted higher-level semantic comprehension rather than single word comprehension (decoding ^21,50^). Additionally, this stimulus set was designed for language levels of individuals >5 years, which could provide seamless comparability across development, and the predictability of each sentence was measured via CP. Furthermore, this stimulus set elicited the LPC (P600), which was reportedly different in individuals with ASD ^33,44,45^. Future use of this stimulus set may increase the comparability and reproducibility within auditory/lexical ASD semantic processing research.

In conclusion, this auditory stimulus set successfully elicited an N400 response as well as other semantically related ERP components: the LPC (P600) and RP. This stimulus set can be used for a wide range of ages and populations, and its availability could contribute to more comparable, consistent results between future studies that use the N400 ERP component to derive information about sentential semantic integration.

## 4. Methods

### 4.1 Participants

Twenty-four neurotypical adults were recruited and provided written informed consent to participate in this study. Four subjects were excluded from data analysis due to failure to remain alert or to sit still during data collection (n=2), or due to noisy EEG data (n=2) (Supplementary Figure 1). The remaining participants make up the fully-analyzed dataset. These twenty subjects ranged in age from 18 to 35 (mean age = 25.5 +/− 4.36), nine were female, and three were left-handed. Every participant spoke English as their first language, and twelve participants were mono-lingual while eight participants reported being bi- or multi-lingual. Demographic information for all participants, including those removed from further analysis are reported in Table 1.

### 4.2 Stimuli

The semantic anomaly paradigm consisted of 221 sentence pairs with incongruent and congruent endings. Nineteen sentence pairs were eliminated before analysis because their endings did not match in syllable number, contained hyphenated phrases, cultural references, or upon closer examination, the supposed incongruent endings were in fact congruent. The remaining 202 sentence pairs ranged from four to eight words in length and were constructed from a list of 300 words selected for high-written word frequency in a child’s everyday environment ^53^. This was to ensure that each sentence could be readily comprehended by typically developing (TD) individuals aged 5 years or older.

This stimulus set included sentence pairs where the incongruent endings were just semantic errors (264 stimuli). These incongruent endings were matched to their congruent pair in word type (e.g. noun or verb) and number (e.g. plural or singular). There were also sentence pairs where incongruent endings contained both semantic and syntactic errors (138 stimuli). Semantically incongruent endings were also classified as syntactic errors if the ending deviated from the syntax provided by the sentential context. The type of syntactic errors included: 1. Endings where a plural noun expectation was deviated with a singular noun or vice versa (54 Stimuli); 2. Endings where an adjective expectation was deviated with a noun or vice versa (40 Stimuli); and 3. Endings where a verb expectation was deviated with a noun or vice versa (44 Stimuli). During final analysis, these 3 types of syntactic errors were combined into a single LD in order to compare the overall response to sentences with both semantic and syntactic errors, and the response to sentences with only semantic errors.

Individual words from the word list were recorded from a female speaker, instructed to voice the words with minimal inflection, stress, and intonation (i.e. in a monotonous non-prosodic manner). Words were then compiled into complete sentences using the Audacity Software (Version 3.0.0. Audacity® software is copyright © 1999 - 2021 Audacity Team. https://audacityteam.org/). These artificially-compiled sentences were manually adjusted to have similar pitch-frequency and pacing between each word within a sentence, and between all sentences.

### 4.3 Procedure

Participants were fit with a 128-electrode cap (Bio Semi B.V. Amsterdam, the Netherlands) and seated in a sound attenuating, electrically shielded booth (Industrial Acoustics Company, The Bronx, NY) with a computer monitor (Acer Predator Z35 Curved HD, Acer Inc.) and a standard keyboard (Dell Inc.). The task was created with Presentation® Software (Version 18.0, Neurobehavioral Systems, Inc. Berkeley, CA). The task was first explained to the participant during the consent process and then again before the experimental session. Individuals were asked to refrain from excessive movement and to focus on a fixation cross throughout the task in order to reduce movement artifacts. The experimental session began by explaining the task for a third time. All instructions were presented both visually on the screen and auditorily through the headphones (Sennheiser electronic GmbH & Co. KG, USA). Instructions were followed by two practice trials which were the same for every participant. Feedback was given about a participant’s response only during practice trials and not during experimental trials. Trials were presented as follows: 1. A fixation cross was on the screen while an auditory sentence stimulus was presented through headphones; 2. A two second pause; and 3. A question (presented both visually and auditorily) asked the participant if the sentence ended as expected, where subjects responded with a right or left arrow key when sentences ended as expected (congruent) or unexpected (incongruent) respectively, to end the trial. A two-second delay was inserted between a subject’s response and the start of the next sentence. During the experiment, a total of 440 stimuli were presented to participants in the same order. This was done to ensure that every participant had the same experience throughout the task for every sentence. Stimuli were separated into 11 blocks with optional breaks between each block. Participants could continue onto the next block by pressing the spacebar.

### 4.4 Data preprocessing

EEG data were preprocessed and analyzed using in-house scripts leveraging EEGLAB functions (Delorme & Makeig, 2004). Data were sampled at a rate of 512Hz, and filtered using a Chebyshev II filter with a band pass of 0.1 - 45 Hz. Prior to analysis, the time from the beginning of a sentence to the onset of the last word was measured for each stimulus using Praat (PRAAT v. 6.1, University of Amsterdam, the Netherlands). These measures were used to adjust the time stamp of each stimulus, so that the data could be aligned to the onset of the last word (i.e. 0 ms), rather than the beginning of the sentence. The epoch was over −200 to 1000 ms relative to the onset of the last word. Epochs were baseline-corrected to the 200 ms interval before the onset of the last word and trials with amplitudes two standard deviations above the mean were rejected. Grand average waveforms were generated by first averaging the trials per condition per electrode, and then averaging by participant.

### 4.5 Statistical analysis

JASP (Jeffrey’s Amazing Statistic Program Team [2020], Version 0.12.2) was used for statistical analyses. Three midline electrodes (Fz, Cz, and Pz) were chosen *a priori* for investigation ^5–7^. Other electrodes (F7, T7, and P7) were investigated *post hoc*. For every participant, these selected electrodes were assessed for an effect of condition using a repeated measure ANOVA (rmANOVA) at four time points of interest (250 ms, 400 ms, 600 ms, and 700 ms). Amplitude values for these electrodes were acquired by averaging the amplitudes across a 10 ms time window, centered at the time point of interest. Additional rmANOVA’s were conducted to assess for a main effect of CP, order, linguistic division, and time. F-scores and p-values for a main effect of condition at the midline electrodes are shown in Table 2. Other main effects for midline electrodes are shown in Table 3. All main effects for electrodes F7, T7, and P7 are shown in Table 4.

Topography plot statistics were generated using the FieldTrip toolbox ^54^ for MATLAB and displayed using the EEGLAB toolbox. A group level cluster-based permutation test was conducted using two-tailed, independent sample t-statistics with a critical alpha-level of 0.05. This test applied the Monte-Carlo method to estimate significance probability, the triangulation method of the neighbours function for spatial clustering, and a multiple-comparison correction. Single sample clusters were combined using maxsum and a 5% two-sided cutoff criterion was applied to both positive and negative clusters. Topography statistics are presented as the average significance over a 10 ms time window centered at the time point of interest. The cluster plot of p-values for all channels were shown in Supplementary Figure 2.

### 4.6 Cloze probability

To further characterize the stimulus set, a RedCap survey was employed to test the CP of all sentence endings. Each sentence in the set was presented with the final word missing (blank) and participants were required to fill in this blank with the first singular word that came to mind. If participants could not think of an answer, they were encouraged to guess rather than leave a blank. Non-answers were not counted towards CP scores; participants were removed from the survey data if they answered fewer than 10 questions out of the 221; and participants were removed if their percent correct was three standard deviations from the mean. After elimination, the survey used the responses from 134 individuals to assess the CP of each sentence. The majority of these stimuli had greater than 80% CP. The CP distribution of sentences were shown in Supplementary Figure 3.

### 4.7 Data Availability

This stimulus set and the supporting datasets are available through Dryad for the scientific community to use freely in their experiments [web link to dataset]. We provide the auditory files for all 442 stimuli, as well as a .tsv file with each stimulus written out for research investigating lexical semantic comprehension. The 24 datasets are provided in BIDS format along with information about the participants and their responses (.tsv and .json). The full dataset includes the raw EEG data (.bdf and .mat) along with the corresponding event, channel rejection, electrode positioning, and filtering information (.tsv and/or .json) for each participant. Additionally, we provide the preprocessed ERP derivatives for this study (.mat) along with the corresponding trial rejection (.tsv) and the filtering and rejection parameters (.json). Information about each stimulus such as stimulus duration, target word onset, group categorization for each division (i.e. CP, order, linguistic error, and time), and the corresponding audio file is provided (.tsv). Lastly, the answers and results of the cloze probability survey are included (.tsv and .json). Refer to the README.txt file in order to access the dataset and use this stimulus set appropriately. Use of this stimulus set or presenting examples from this stimulus set should include a citation to this paper.

### 4.8 Code Availability

The code generated for the analysis of these datasets is available through Zenodo via Dryad [web link to dataset]. The provided code was utilized to create the preprocessed ERP derivatives as well as figure components.

## Supplementary Files

Upon acceptance for publication, the authors will coordinate with the editorial office to ensure that the entire stimulus set and respective analysis documents, as well as the full and appropriately de-identified datasets are uploaded to a publicly available repository and that the appropriate link is made to the article file.

## Ethics and Consent

The Research Subjects Review Board of the University of Rochester approved all the experimental procedures (STUDY00002036). Each participant provided written informed consent in accordance with the tenets laid out in the Declaration of Helsinki.

## Acknowledgements

The authors acknowledge the contribution of Mr. Oren Bazer for assistance with data collection, and Dr. Kathryn-Mary Wakim and Mr. David Richardson for their contributions to the analysis pipeline.

## Authors’ Contributions

All authors collectively conceived of the study. KKT created the stimulus set, recruited participants, and collected the data. KKT analyzed the data in consultation with EGF and JJF, and produced the figures for this study. KKT wrote the first draft of the manuscript and all authors provided editorial input on subsequent drafts. All authors read and approved the final version of this manuscript.

## Supplemental Figures

**Supplementary Figure 1:**
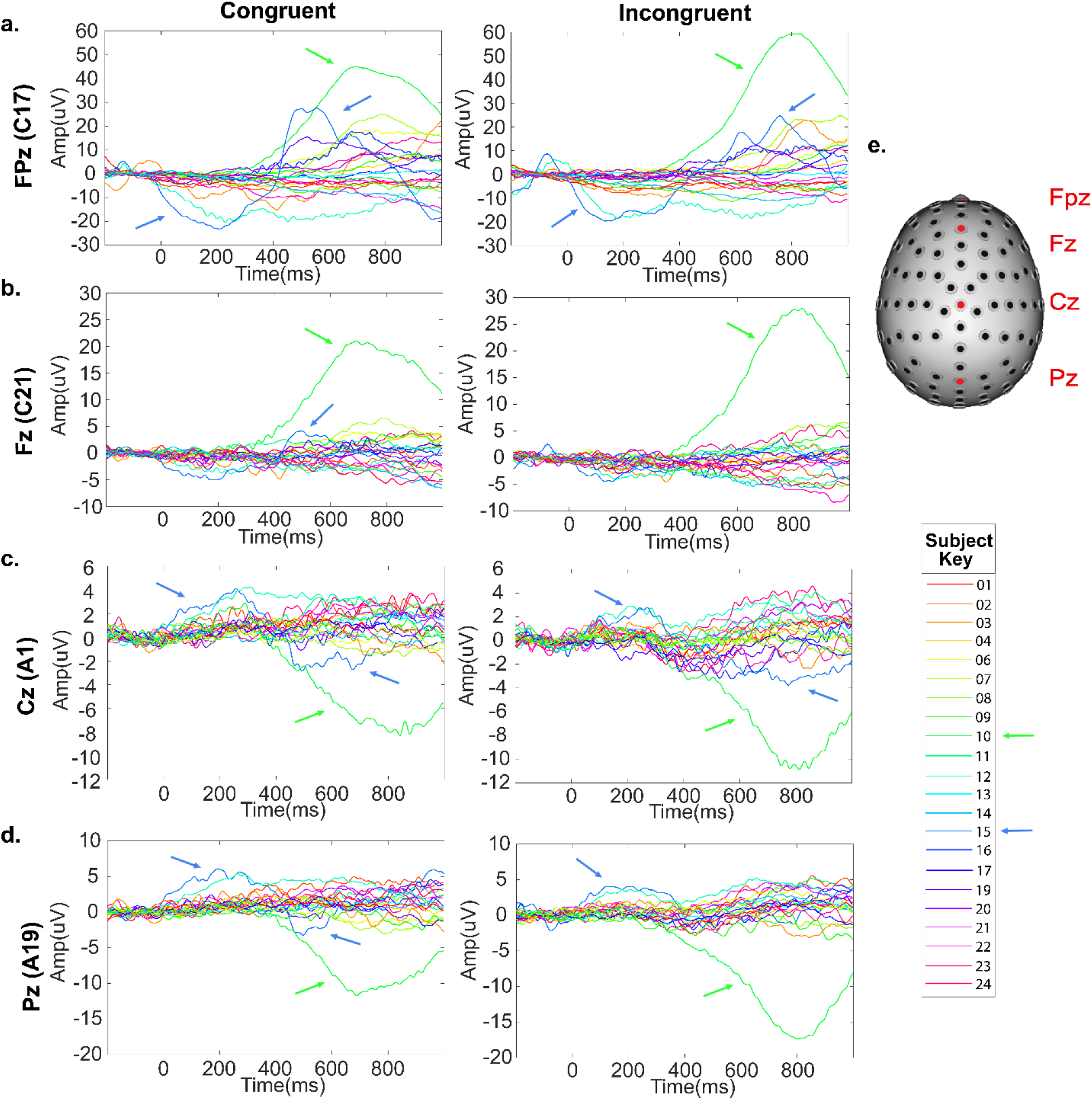
ERP plots supporting the rejection of subject 10 and 15 from final analysis. ERP plots for midline Electrodes **a.** FPz, **b.** Fz, **c.** Cz, and **d.** Pz, separated by subject for both congruent and incongruent conditions, using all 440 stimuli. Arrows are pointing to the EEG activity of subject 10 (Green) and subject 15 (Blue). **e.** Head model of electrode locations and subject key. Both subject 10 and 15 displayed deflections that we could not attribute to brain activity. Subject 10 was eliminated for drastically large amplitudes between 400 and 1000 ms in nearly all 128 electrodes, but primarily frontal electrodes. Subject 15 was eliminated due to an atypical waveform consisting of a large early deflection between 0 ms and 400 ms followed by a reverse deflection between 400 ms and 800 ms. This was seen in all fontal electrodes, but was significantly more pronounced in the most exterior frontal electrodes (Supplementary Figure 1a).

**Supplementary Figure 2:**
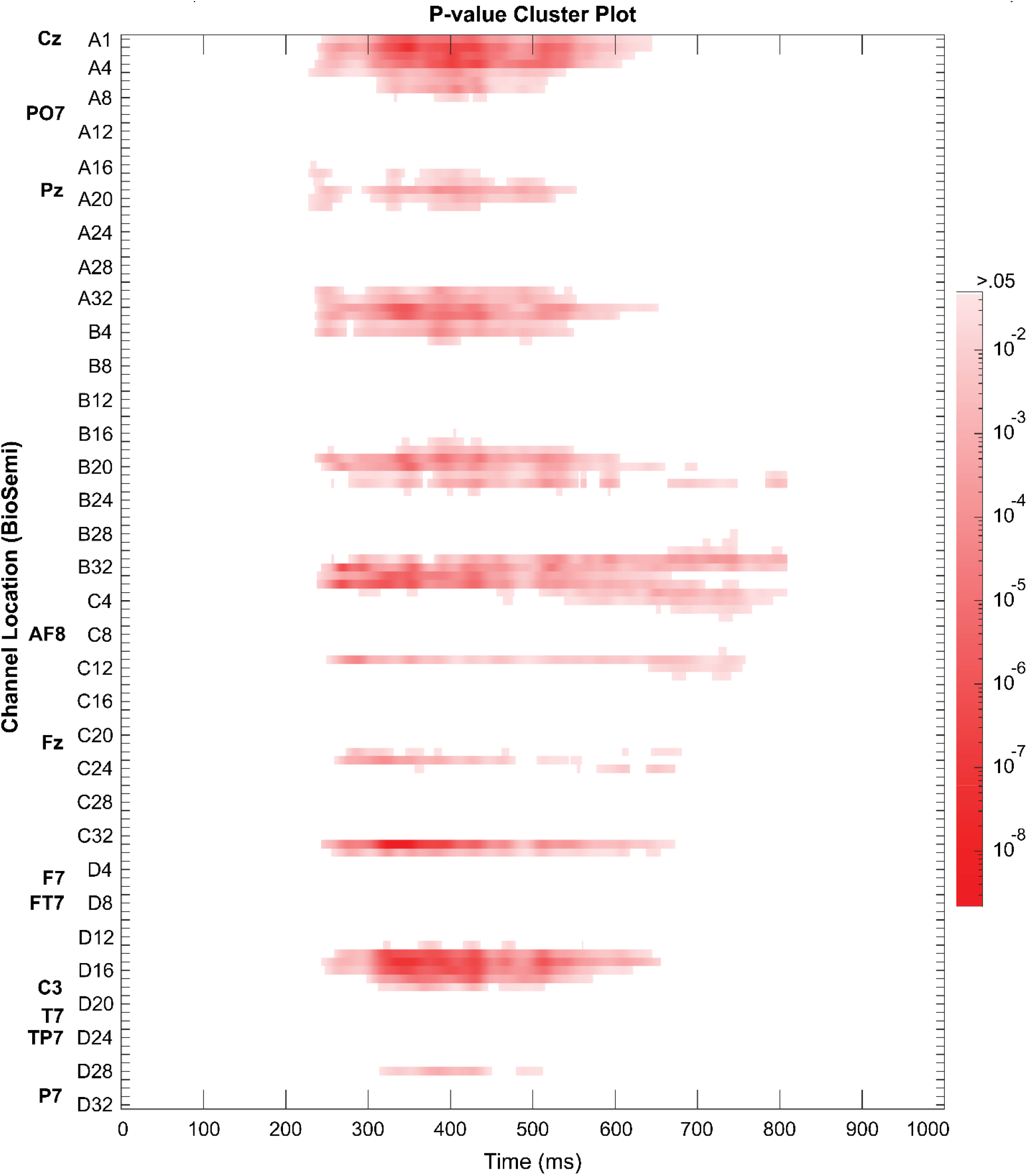
Cluster plot of p-values for all 128 channels after conducting two-tailed independent sample t-tests. Red is significant while white means that p-values >.05. T-tests used the Monte-Carlo method, the triangulation method for spatial clustering, and a multiple-comparison correction. A 5% two-sided cutoff criterion was applied to both positive and negative clusters. Channels are identified by BioSemi cap coordinates and the electrodes mentioned in this dataset are noted with the standard 10 - 20 electrode positions.

**Supplementary Figure 3:**
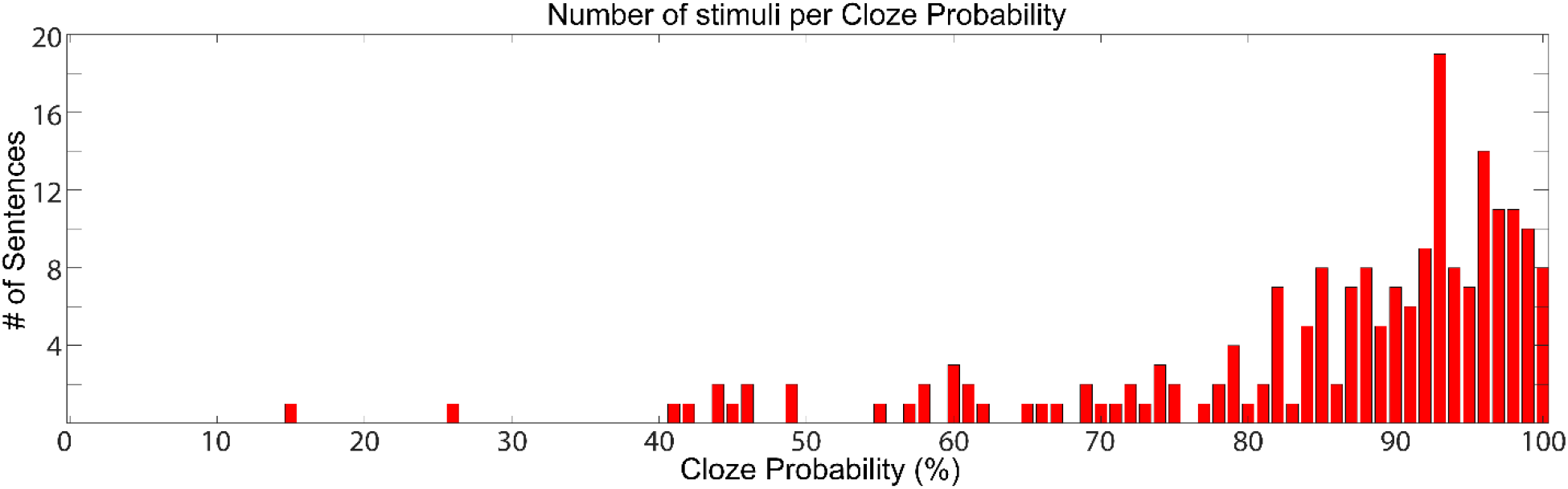
Bar graph showing the distribution of Cloze Probability for the congruent stimuli analyzed in this dataset. Total congruent stimuli assessed, 202 sentences.

**Supplementary Figure 4:**
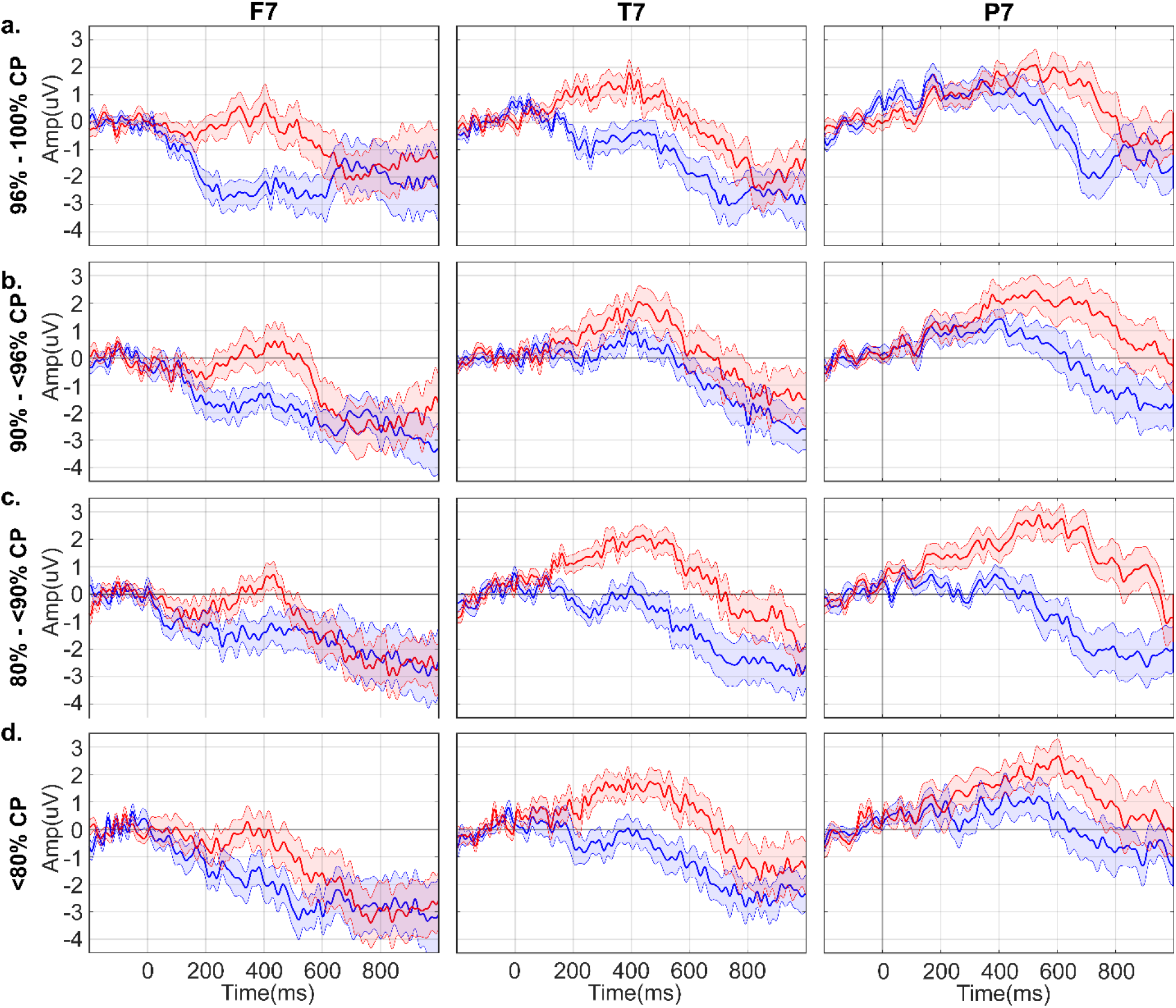
ERP plots for left frontal (F7), temporal (T7), and temporal-occipital (P7) electrodes separated by differing levels of Cloze Probability. The ERP plots show amplitude changes over 1000 ms, per condition (Blue, congruent. Red, Incongruent). The shading around each line represents the s.e.m. **a.** Sentences with 96 - 100% cloze probability (54 congruent and 55 incongruent stimuli). **b.** Sentences with 90 - <96% cloze probability (57 sentences pairs). **c.** Sentences with 80 - <90% cloze probability (47 sentence pairs). **d.** Sentences with <80% cloze probability (43 congruent and 42 incongruent stimuli). Statistics are on Table 4.

**Supplementary Figure 5:**
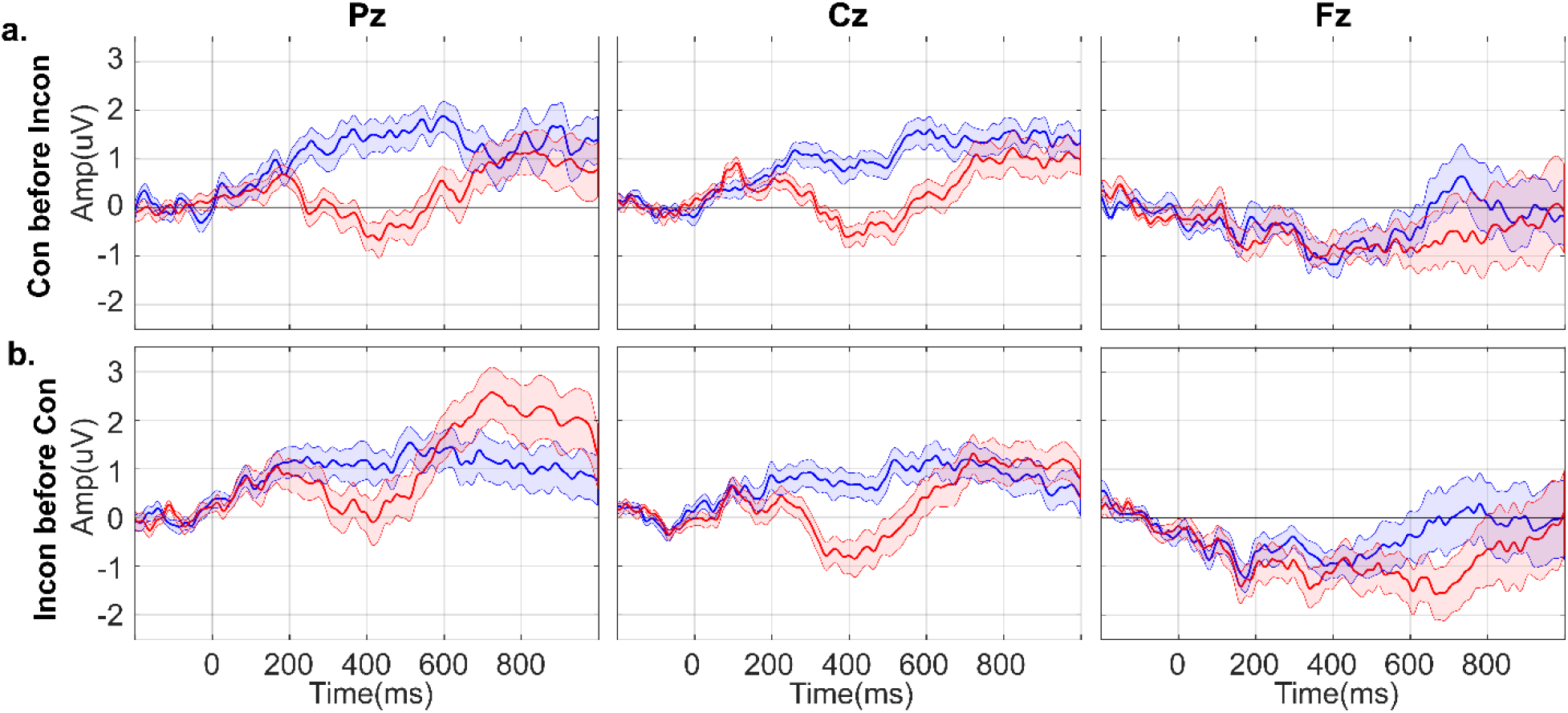
ERP plots for midline electrodes Pz, Cz, and Fz, separated by the order of stimulus presentation during the experiment. Activity over 1000 ms per condition (Blue, congruent. Red, Incongruent). The shading around each line represents the s.e.m. **a.** Congruent sentence pairs presented prior to their corresponding incongruent ending sentence pair (99 congruent and 100 incongruent stimuli). **b.** Incongruent sentence pairs presented prior to their corresponding congruent ending sentence pair (102 congruent and 101 incongruent stimuli). Statistics are on Table 3.

**Supplementary Figure 6:**
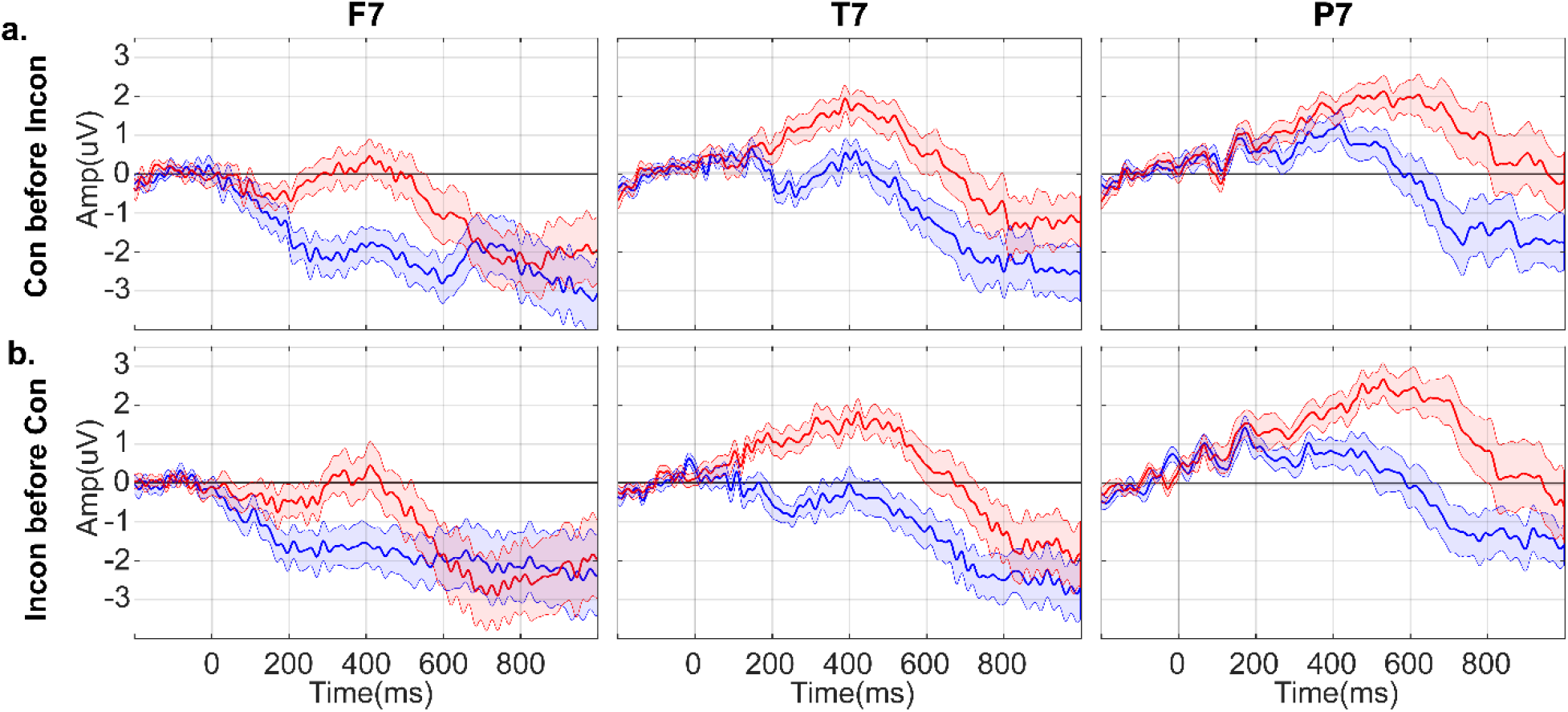
ERP plots for left frontal (F7), temporal (T7), and temporo-occipital (P7) electrodes, separated by the order of stimulus presentation during the experiment. Activity over 1000 ms per condition (Blue, congruent. Red, Incongruent). The shading around each line represents the s.e.m. **a.** Congruent sentence pairs presented prior to their corresponding incongruent ending sentence pair (99 congruent and 100 incongruent stimuli). **b.** Incongruent sentence pairs presented prior to their corresponding congruent ending sentence pair (102 congruent and 101 incongruent stimuli). Statistics are on Table 4.

**Supplementary Figure 7:**
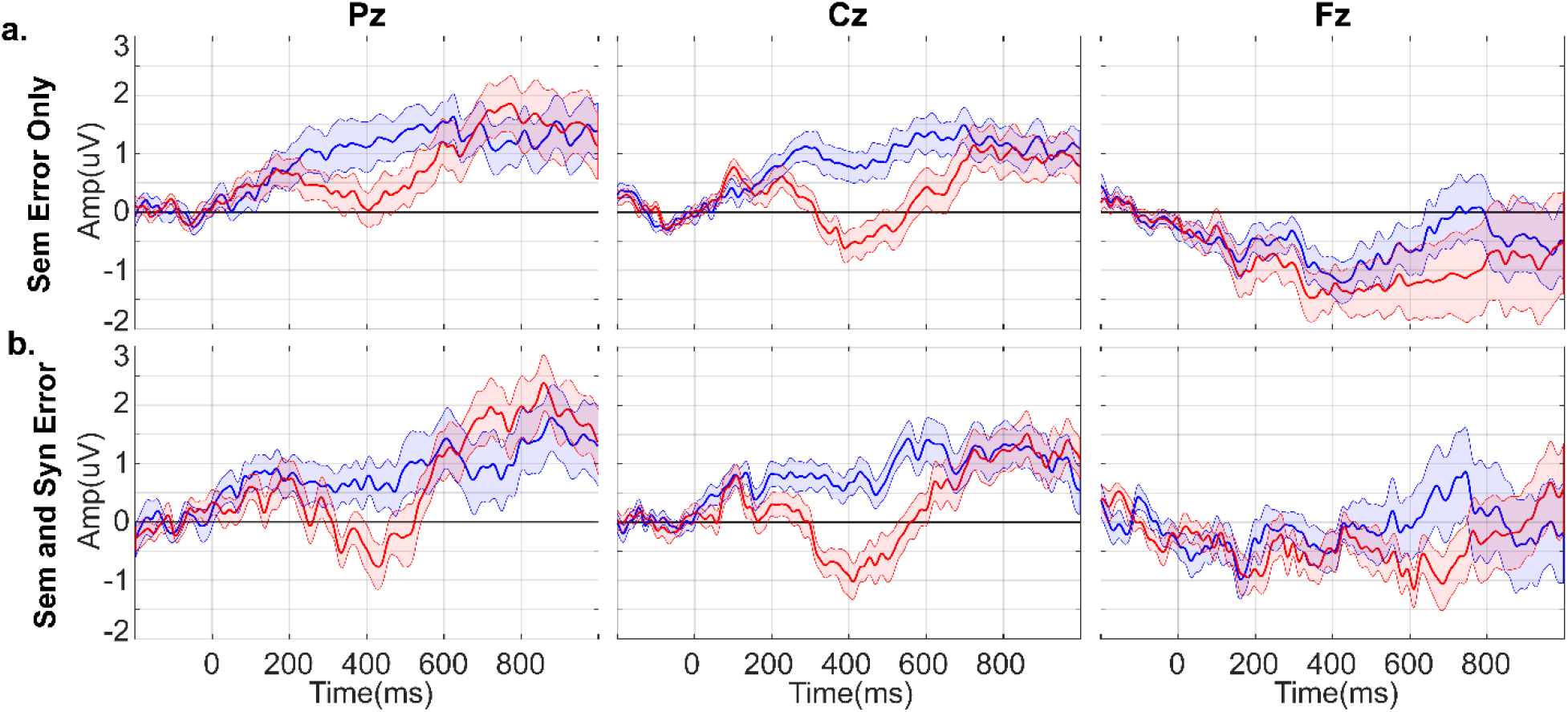
ERP plot for midline electrodes Pz, Cz, and Fz, separated by linguistic ending errors. Activity over 1000 ms per condition (Blue, congruent. Red, Incongruent). The shading around each line represents the s.e.m. **a.** Incongruent sentence stimuli with only semantic ending errors and their congruent pair (132 sentence pairs) **b.** Incongruent sentence stimuli with both semantic and syntactic ending errors and their congruent pair (69 sentence pairs).Statistics are on Table 3.

**Supplementary Figure 8:**
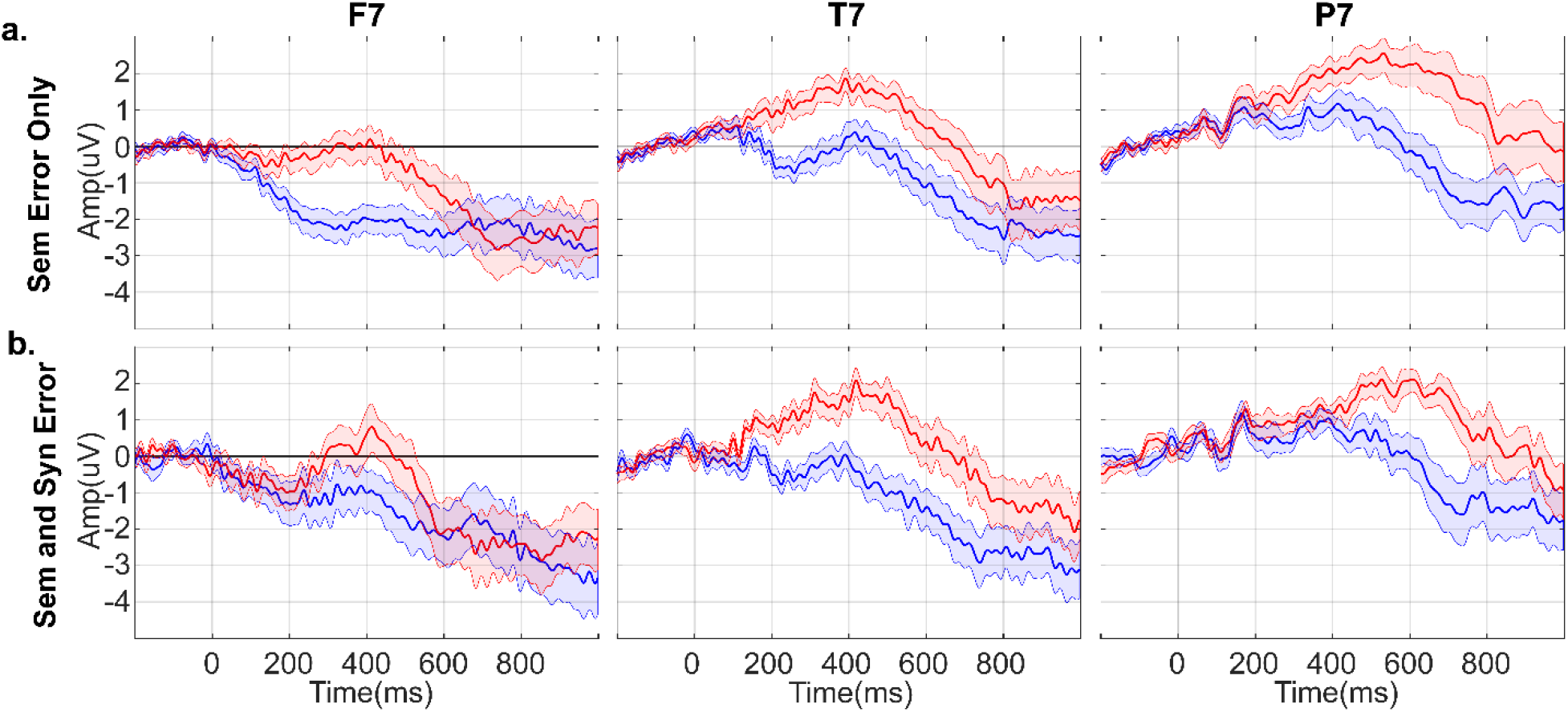
ERP plots for left frontal (F7), temporal (T7), and temporo-occipital (P7) electrodes, separated by linguistic ending errors. Activity over 1000 ms per condition (Blue, congruent. Red, Incongruent). The shading around each line represents the s.e.m. **a.** Incongruent sentence stimuli with only semantic ending errors and their congruent pair (132 sentence pairs) **b.** Incongruent sentence stimuli with both semantic and syntactic ending errors and their congruent pair (69 sentence pairs). Statistics are on Table 4.

**Supplementary Figure 9:**
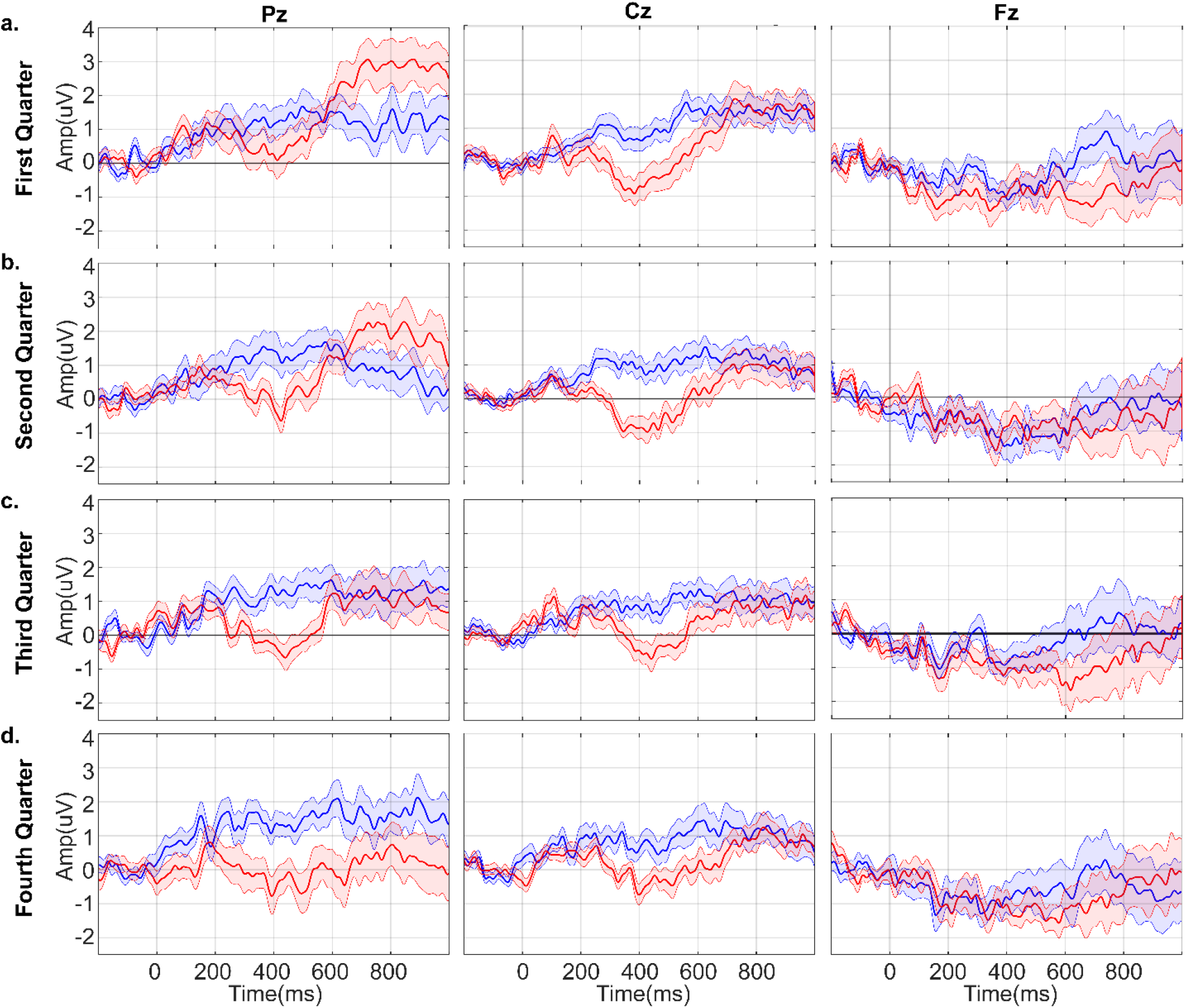
ERP plots for midline electrodes Pz, Cz, and Fz, separated by quarters of the experiment. Activity over 1000 ms per condition (Blue, congruent. Red, Incongruent). The shading around each line represents the s.e.m. **a.** First quarter of the experiment (51 congruent and 48 incongruent stimuli) **b.** Second quarter of the experiment (46 congruent and 53 incongruent stimuli). **c.** Third quarter of the experiment (52 congruent and 50 incongruent stimuli). **d.** Last quarter of the experiment (52 congruent and 50 incongruent stimuli). Statistics are on Table 3.

**Supplementary Figure 10:**
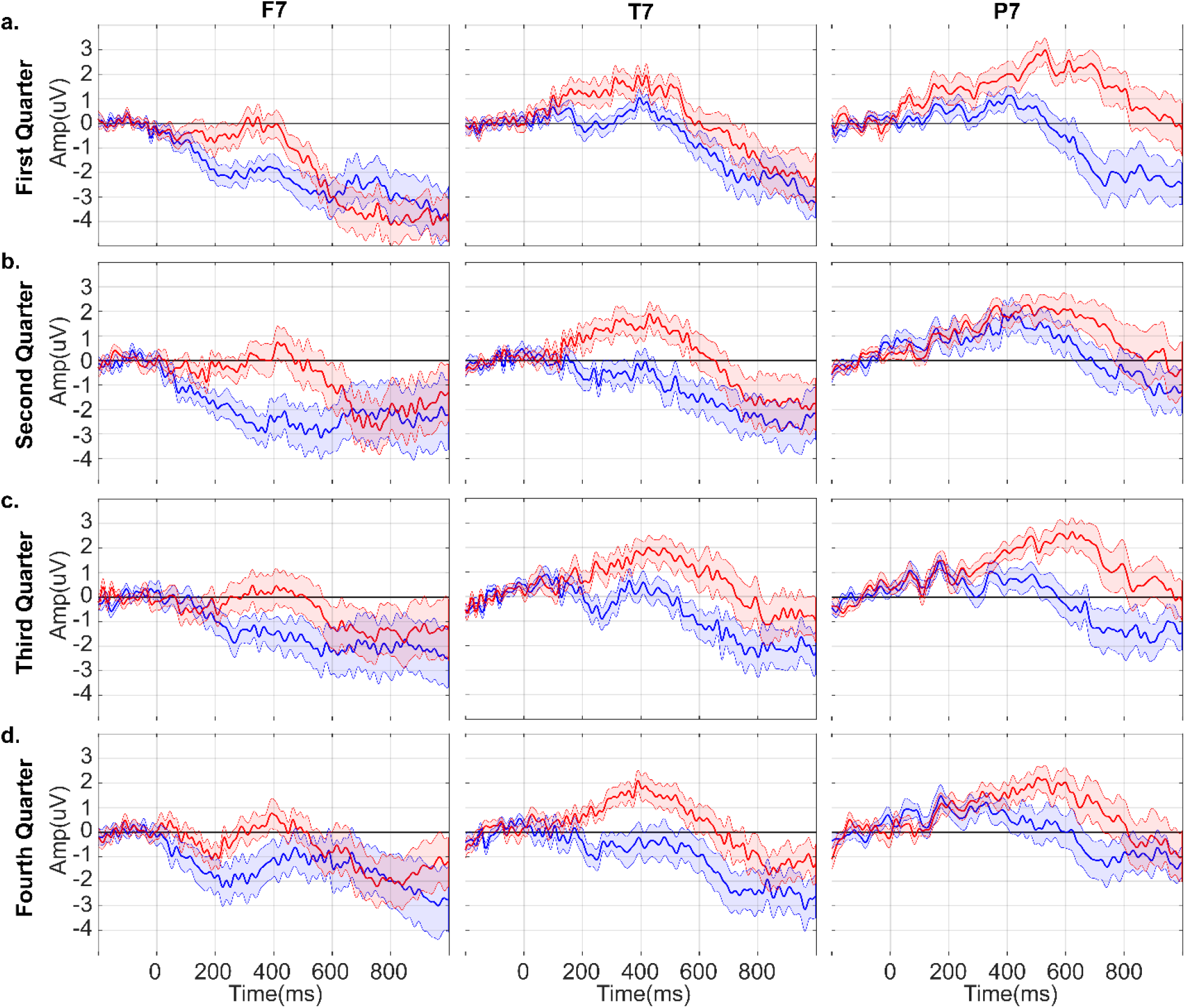
ERP plots for left frontal (F7), temporal (T7), and temporo-occipital (P7) electrodes, separated by quarters of the experiment. Activity over 1000 ms per condition (Blue, congruent. Red, Incongruent). The shading around each line represents the s.e.m. **a.** First quarter of the experiment (51 congruent and 48 incongruent stimuli) **b.** Second quarter of the experiment (46 congruent and 53 incongruent stimuli). **c.** Third quarter of the experiment (52 congruent and 50 incongruent stimuli). **d.** Last quarter of the experiment (52 congruent and 50 incongruent stimuli). Statistics are on Table 4.

